# Breast cancer cell-derived extracellular vesicles accelerate collagen fibrillogenesis and integrate into the matrix

**DOI:** 10.1101/2024.08.08.607183

**Authors:** Nicky W. Tam, Rumiana Dimova, Amaia Cipitria

## Abstract

Extracellular vesicles (EVs) within the extracellular matrix (ECM) are often studied as passive elements whose diffusion and behaviour are subject to the composition and structure of the ECM. While EV diffusion and distribution in tissues are indeed governed by matrix interactions, accumulating evidence suggests that EVs contain much of the cellular machinery required for actively remodeling ECM. Using rheology and confocal reflectance microscopy, we investigate the gelation of collagen I hydrogels formed in the presence of EVs, and show that EVs can play an active role in ECM formation. EVs appear to nucleate new fibrils, recruiting collagen molecules from solution and accelerating their polymerization. Trypsinization of EVs shows that collagen-EV interactions are primarily mediated by surface proteins. The use of extruded plasma membrane vesicles shows that membrane composition determines final fibril length and matrix structure. EVs also become integrated into the fibril structures that they help form, reminiscent of matrix vesicles found in situ within tissues. This represents a plausible way by which EVs are deposited into the ECM, becoming signaling cues for resident cells. Our data show that EV-matrix interactions are dynamic and can contribute to the remodeling of tissue microenvironments.

**Significance:** Extracellular vesicles (EVs) are nanoscale membrane structures known for their role in facilitating cellto-cell trafficking of proteins, lipids, RNA, and other signaling molecules. In this report, we show that EVs derived from breast cancer cells are not merely passive messengers, but also direct active effectors of extracellular matrix (ECM) remodeling processes. Bulk rheology and confocal microscopy show that these EVs have the ability to nucleate new collagen fibrils and accelerate the formation of dense fibrillar collagenous networks. This has important implications in cancer pathology, where matrix density is often associated with worse disease outcomes, but could also potentially be exploitable in future tissue engineering applications.

## Introduction

Dynamic, reciprocal interactions between cells and the extracellular matrix (ECM) help to shape tissue structure and function. Extracellular vesicles (EVs) are one component of a cell’s arsenal of signaling modalities (1, 2) and are mostly appreciated for their role as cell-to-cell messengers of signaling cues, influencing the tissue microenvironment through cell proxies (3–5). This is achieved in a number of ways, through promoting the proliferation of ECM-secreting cells (6), changing the ECM secretion profile of cells (7– 10), or increasing the expression of ECM-degrading matrix metalloproteinase enzymes (MMPs) (11, 12). Mounting evidence suggests, however, that EVs themselves may play a direct role in remodeling the ECM. Crosslinking enzymes, such as transglutaminase (11) or lysyl oxidase (12), as well as lytic enzymes, including MMPs (13, 14) and heparinase (15) have been found in EVs, allowing them to act as direct effectors of remodeling processes. In this report, we investigate EV interactions with collagen I to better understand how EVs may play a role in the formation of collagen fibril structures.

Collagen I constitutes one of the most abundant proteins in mammalian tissues and plays important roles both structurally and in terms of cell signaling within tissues (16–18). The synthesis, secretion, and assembly of collagen I into fibrillar structures is known as fibrillogenesis and is a tightly regulated phenomenon involving many different binding partners and molecular processes *in vivo* (19, 20). The functional fibril-forming unit of collagen I is tropocollagen, itself a three-stranded helix composed of individual monomeric chains. These monomers can be recovered and separated with SDS-PAGE from samples of solubilized collagen I obtained through enzymatic or acid extraction of tissues, as is done commercially (21–23). It is not immediately clear, however, if molecules in solution exist in this way as free monomers or as triple-helical tropocollagen units. Nevertheless, collagen fibrillogenesis *in vitro* represents the spontaneous aggregation of tropocollagen bundles, followed by end-to-end and lateral joining of nucleated bundles into fibrils (24–26). While this process may not fully replicate the predominantly fibroblast-mediated deposition of collagenous ECM (20, 27, 28), understanding how EVs might influence the kinetics of this baseline spontaneous fibril formation can provide important insight into how EVs contribute to the dynamic nature of tissue microenvironments.

Previously, we and others have reported that EVs can interact with collagen I *via* integrin receptors and that these interactions modulate their ability to diffuse and infiltrate into reconstituted collagen hydrogels (29, 30). Here, we explore how EVs may in turn affect collagen fibrillogenesis and how they may play an active role in ECM remodeling processes. To this end, we employ bulk rheology to investigate how EVs affect the gelation kinetics and fibril formation of collagen I. We also employ confocal reflectance microscopy, fluorescence confocal microscopy, and image analysis techniques to investigate how EVs affect collagen fibril structure, as well as how EV-fibril interactions affect localization of EVs in a fibril matrix. By comparing the behaviour of EVs with extruded plasma membrane vesicles (30, 31) and synthetic liposomes, we show that EVs have a distinct effect on collagen fibril formation that suggests a specialized biological role in nucleating and accelerating fibrillogenesis.

## Results and Discussion

### Generation and collection of breast cancer cell-derived EVs

EVs were generated by MDA-MB-231 breast cancer cells cultured in serum-free medium to avoid contamination from serum-derived vesicles. Collection and purification of EVs was conducted, as previously published, using size exclusion chromatography (SEC) (32–34). We have previously characterized EVs collected and purified in this manner with cryogenic scanning electron microscopy and Western blot analysis of common EV protein markers, showing that such EVs largely consist of 100-400 nm diameter exosomes of endosomal origin with high enrichment of integrin β1 (ITGB1) and CD63 (30). Analysis of particle sizes in SEC fractions with dynamic light scattering (DLS) can be found in the Supporting Materials (Supplemental Figure S1).

The MDA-MB-231 cell line was chosen as it is a commonly used model in breast cancer research for highly-malignant triple-negative type breast cancers (35). EVs generated from this cell line have previously been used in other studies (30, 36, 37), showing that they are a reliable source of EV material.

### Breast cancer cell-derived EVs accelerate collagen I fibrillogenesis

To begin to probe the interactions between EVs and collagen I, we used bulk rheology in oscillatory shear mode to assess how the presence of EVs affects the gelation kinetics of collagen I hydrogels. Two different buffers were used for experiments to also investigate calcium-dependence with respect to integrin binding: a calcium-free HEPES-buffered saline (HBS, pH 7) and a HEPES-buffered saline supplemented with 2mM calcium chloride (HBS+Ca, pH 7). Figure 1A,B shows a general schematic of how the rheology was conducted and analyzed. Collagen solutions with EVs were mixed directly on the stage of a rheometer, cooled below 7°C to prevent premature gelation. Once mixed, the stage was heated to 35 °C over approximately 60 s to start the gelation process. The storage (G’) and loss (G”) moduli were measured over time and both were found to exhibit sigmoidal shapes over the gelation process (Fig. 1B), in agreement with the literature (38, 39). While it should theoretically be possible to observe a gel-sol transition, whereby the trajectories of G’ and G” ‘crossover’ to mark the transition from liquid to gel states (38), this was not observed in our data, likely because the transition occurs at lower values of G’ and G” that our rheometer is not sensitive enough to detect.

**Figure 1.**
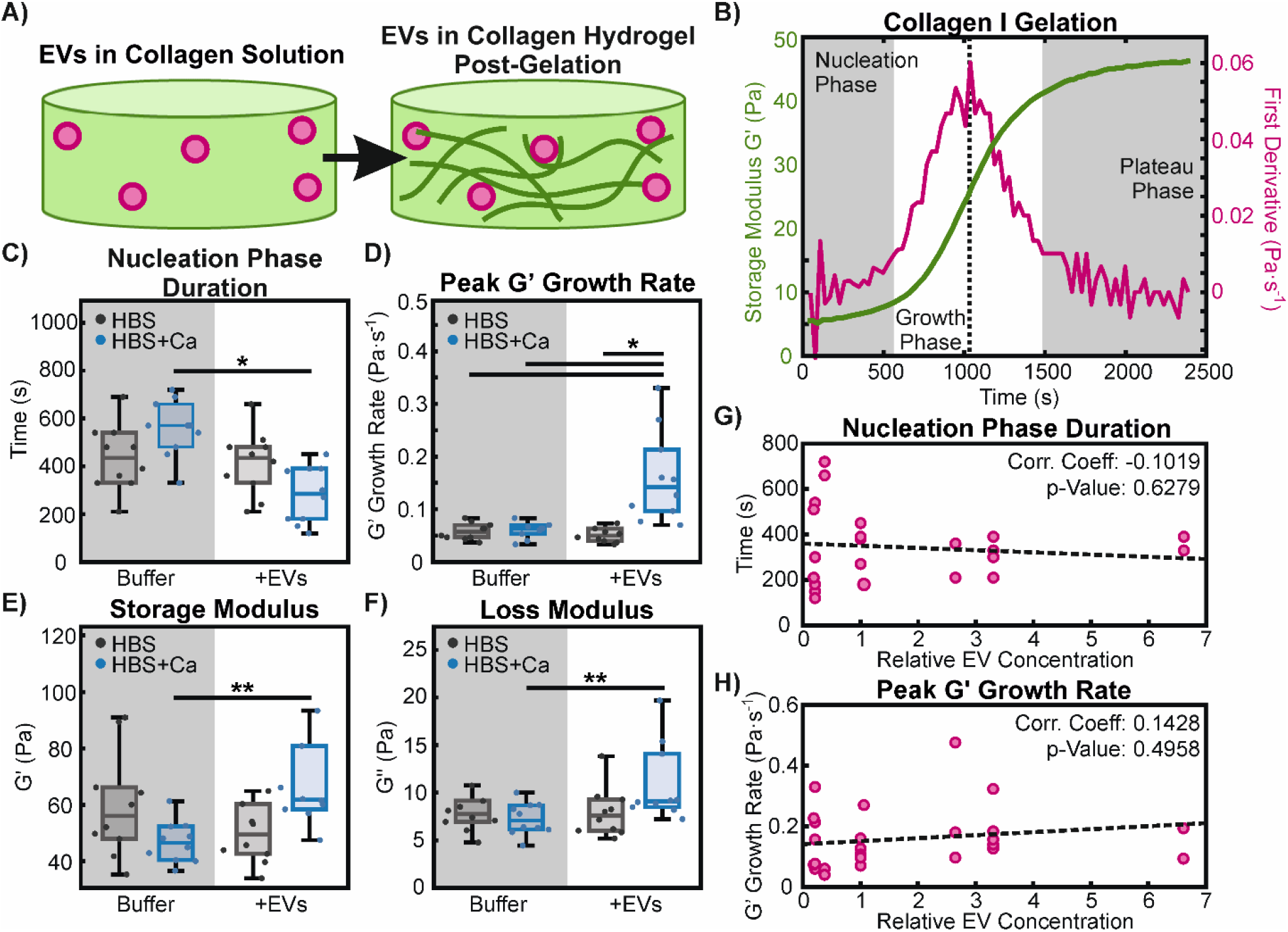
Gelation kinetics of collagen I hydrogels. A) Depiction of EVs present in solution prior to collagen I gelation and the formation of collagen fibrils as gelation progresses. B) Representative example of a collagen gelation curve with calcium but no EVs present. The storage modulus (green) displays a sigmoidal shape and is plotted with its first derivative (magenta). The gelation process is divided into three phases, shown as shaded regions: the nucleation phase, defined as the time until the first derivative exceeds the mean plus two standard deviations of the initial 5 time points used as a baseline; the growth phase; and the plateau phase. A dotted line indicates the inflection point, at which the peak growth rate of G’ is taken. C,D) Comparison of the duration of the nucleation phase (C) and the peak growth rate of G’ (D), taken from gelation curves of hydrogels formed without (Buffer) and with (+EVs) EVs, in calcium-free (HBS; grey) and calcium-containing (HBS+Ca; blue) buffers. Statistically significant differences were determined with 2-way ANOVA with Tukey-Kramer post-hoc analysis, as indicated with * (p<0.05) over n=10 replicates. E,F) Endpoint measurements of storage (E) and loss (F) moduli, measured 1 hour after the start of gelation. A 2-way ANOVA did not show any statistical differences between experimental conditions, but significant differences were found with a 1-way ANOVA with Tukey-Kramer post-hoc analysis over n=10 replicates, as indicated by ** (p<0.05). G,H) Dose response of EVs on collagen I gelation kinetics. EV concentrations are normalized to the average concentration of particles used in all other experiments. Dotted lines show linear regression trends between particle concentrations and nucleation phase duration (G) and peak G’ growth rate (H). Correlation coefficient values are shown, along with calculated p-values, which show a lack of statistically significant dose-dependence (p>0.05) for both the nucleation phase duration and the peak G’ growth rate.

For clarity, and to avoid the greater impact of noise at low amplitudes, we focused on the storage modulus and broke down the gelation process into three distinct phases: (i) an initial lag phase that roughly corresponds to nucleation of collagen aggregates, as described in previous reports (24, 26, 38); (ii) a growth phase, during which the extension of nucleated aggregates into fibril structures rapidly increases the elastic strength of the bulk material; and (iii) a plateau phase, during which the final structure and elastic strength of the gel become set (Fig. 1B). We also determined the peak growth rate of G’ during the growth phase, which occurs at the midpoint or inflection point of the gelation curve. While other parameters, such as the time point at which the inflection point occurs or the start of the plateau phase were analyzed, the duration of the nucleation phase and the peak growth rate appeared to be most affected by the presence of EVs (Fig. 1C,D). In particular, EVs appear to significantly decrease the length of the nucleation phase and increase the peak growth rate of collagen I gelation, but only in the presence of calcium in solution. We also measured endpoint rheology, *i*.*e*. the final storage and loss moduli at the end of the plateau phase when the gel is fully set (Fig. 1E,F). Here, EVs increase the storage and loss moduli, but again, only in the presence of calcium in the buffer. No effect is observed without calcium, strongly suggesting the existence of calcium-dependent machinery, such as integrins, for example, that are involved in the interactions between EVs and collagen I.

To determine if the effects of EVs on collagen gelation are dose-dependent, we varied the amount of EVs used in gelation experiments with calcium-containing buffer, but found no clear trend in the nucleation phase duration, nor the peak growth rate in G’ (Fig. 1G,H). It is possible that all our experiments were conducted at too low an EV concentration range, or possibly in a saturation regime, such that differences in gelation kinetics may vary at lower or higher EV concentrations. Alternatively, differences may occur at the single-fibril level that cannot be easily detected with bulk rheology. For reference, our default normalized particle concentration of 1 in Figure 1G,H corresponds to an in-gel concentration of 2.6±0.4 ×10^7^ particles×µL^-1^. Other studies involving plasma-derived EVs from diabetic patients (40), EVs extracted from human renal cancer tissue (41), and EVs extracted from mouse brain tissue (42) report EV concentrations of up to 1.5×10^9^ particles×µL^-1^, 2.5×10^7^ particles×µL^-1^, and 1.2×10^5^ particles×µL^-1^, respectively.

### Effects on collagen I gelation are particle-specific

To determine whether the effects of EVs on collagen I fibrillogenesis and hydrogel gelation were due to specific molecular interactions or non-specific interactions from simply having a particle inclusion in the gel, we repeated rheology experiments with synthetic large unilamellar vesicles (LUVs) composed of pure 1,2-dioleoyl-snglycero-3-phosphocholine (DOPC), produced as previously described (43). Furthermore, since EVs are known to have a membrane composition distinct from that of their original source cell’s plasma membrane, we wanted to study the importance of membrane composition on interactions with collagen I. We thus tested the effects of extruded large plasma membrane vesicles (LPMVs) (30). These vesicles are formed by extruding plasma membrane material obtained from cells through the use of vesiculation agents (31, 44) and are approximately the same size as our collected EVs while being more representative of whole plasma membrane in terms of composition (Fig. 2A; Suppl. Fig. S1B). Finally, to determine what kind of molecules in EVs are primarily responsible for interactions with collagen I, we treated EVs with trypsin to digest their surface proteins (tEVs). This would knock out protein interactions and possibly also disrupt the glycocalyx to a degree, since glycosylated proteins will be affected. Lipids and other sugar groups, however, would remain largely intact.

**Figure 2.**
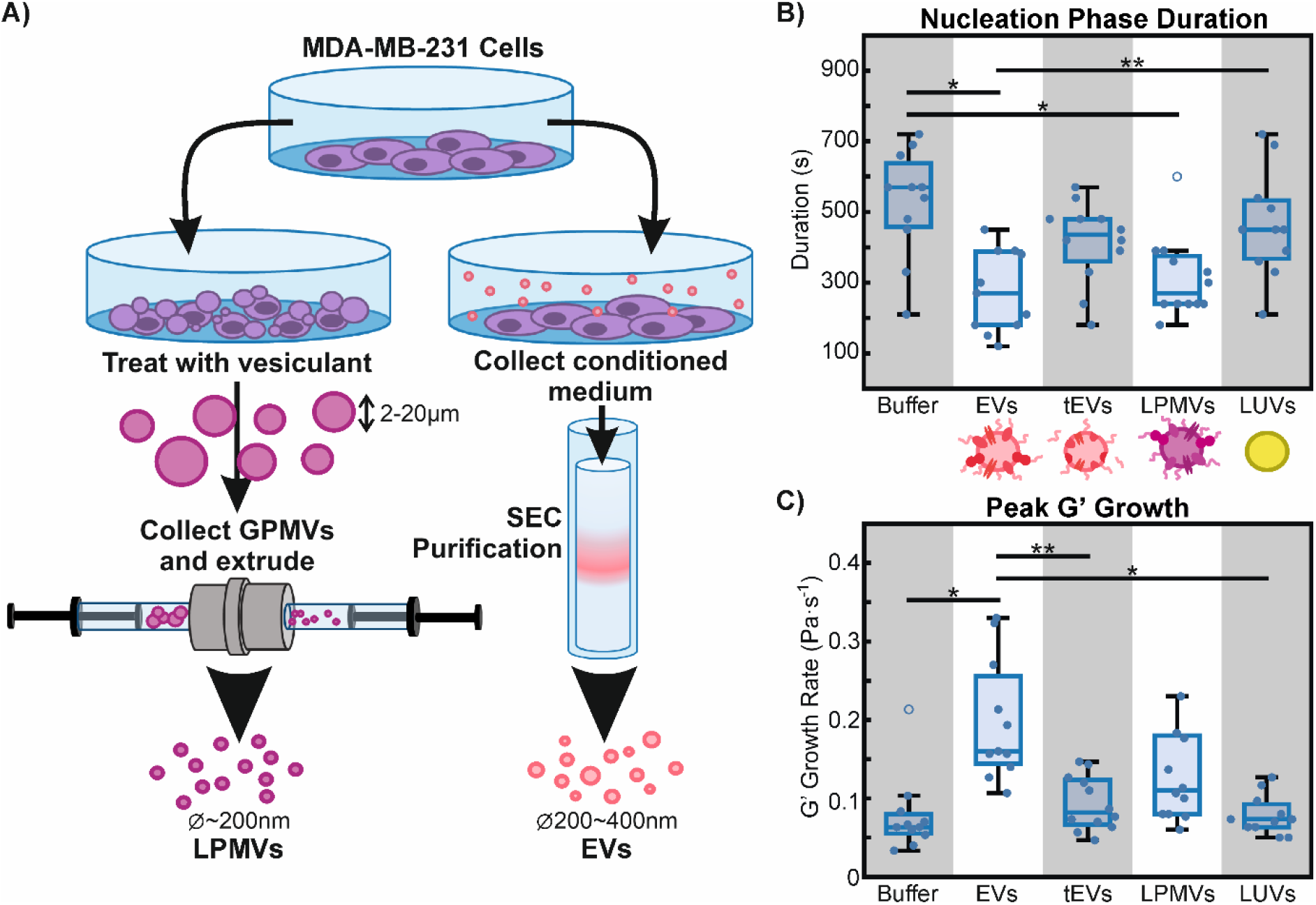
Particle-specific effects on collagen I gelation in the presence of calcium. A) Schematic diagram comparing the production of LPMVs (left) and collection and purification of EVs (right). B) Nucleation phase duration of collagen I gels formed with EVs, trypsinized EVs (tEVs), LPMVs, and synthetic LUVs compared to particle-free controls (Buffer). Cartoons underneath illustrate the differences in the particles in terms of their composition and surface characteristics. C) Peak growth rate of G’ for gels formed without particles (Buffer) and with EVs, tEVs, LPMVs, and LUVs. Statistically significant differences are indicated by black bars, as determined by 1-way ANOVA with Tukey-Kramer post-hoc analysis. Significance levels are indicated by * (p<0.01) and ** (p<0.05). Individual points represent separate experimental replicates (n=12 for tEVs, n=11 for other conditions) while empty circles represent outlier data points that have been excluded from statistical analyses.

We found that nucleation phase duration is significantly reduced by EVs and LPMVs, but not by tEVs or LUVs when compared to the particle-free controls. This suggests that EVs and LPMVs share similar machinery that is lost in tEVs and not present in LUVs, which allows interaction with collagen I. With regards to the peak G’ growth rate, only EVs appear to have a significant effect compared to the particle-free control. LPMVs appear to slightly increase the peak G’ growth rate, but not to a statistically significant extent.

Taken together, this shows that the effects of EVs on collagen I gelation are particle-specific. EVs and LPMVs significantly reduce the nucleation phase duration and EVs alone significantly increase the peak G’ growth rate. Moreover, since tEVs behave similarly to synthetic LUVs and have no effect compared to the particle-free control, it appears that proteins are primarily responsible for the interactions between EVs and collagen I. LPMVs also have intact membrane proteins that can mediate interactions with collagen I, as seen in the decrease in nucleation phase duration. They do not behave exactly the same as EVs, however, as they do not raise the peak G’ growth rate to the same level as EVs. Such functional differences reflect the compositional differences between EVs and LPMVs.

### Breast cancer cell-derived EVs affect collagen I fibril structure and matrix geometry

Confocal reflectance microscopy was used to image the fibrils making up the structure of collagen I hydrogels (41, 47). Confocal stacks were imaged, processed, and analyzed to measure overall fibril content, hydrogel mesh size, and average fibril length (Fig. 3A-E). Fibril content was determined by summing up the number of ‘fibril pixels’ and dividing by the total number of pixels in the image. Hydrogel mesh size was determined with a previously described protocol (41, 47) by determining the sizes and number of spaces between the fibrils in the xand y-directions. These data fall into exponential distributions, from which the mesh sizes were obtained by determining the mathematical mean values of fitted exponential functions. To determine average fibril length, we used the CT-FIRE analysis software developed by Eliceiri et al. (48).

**Figure 3.**
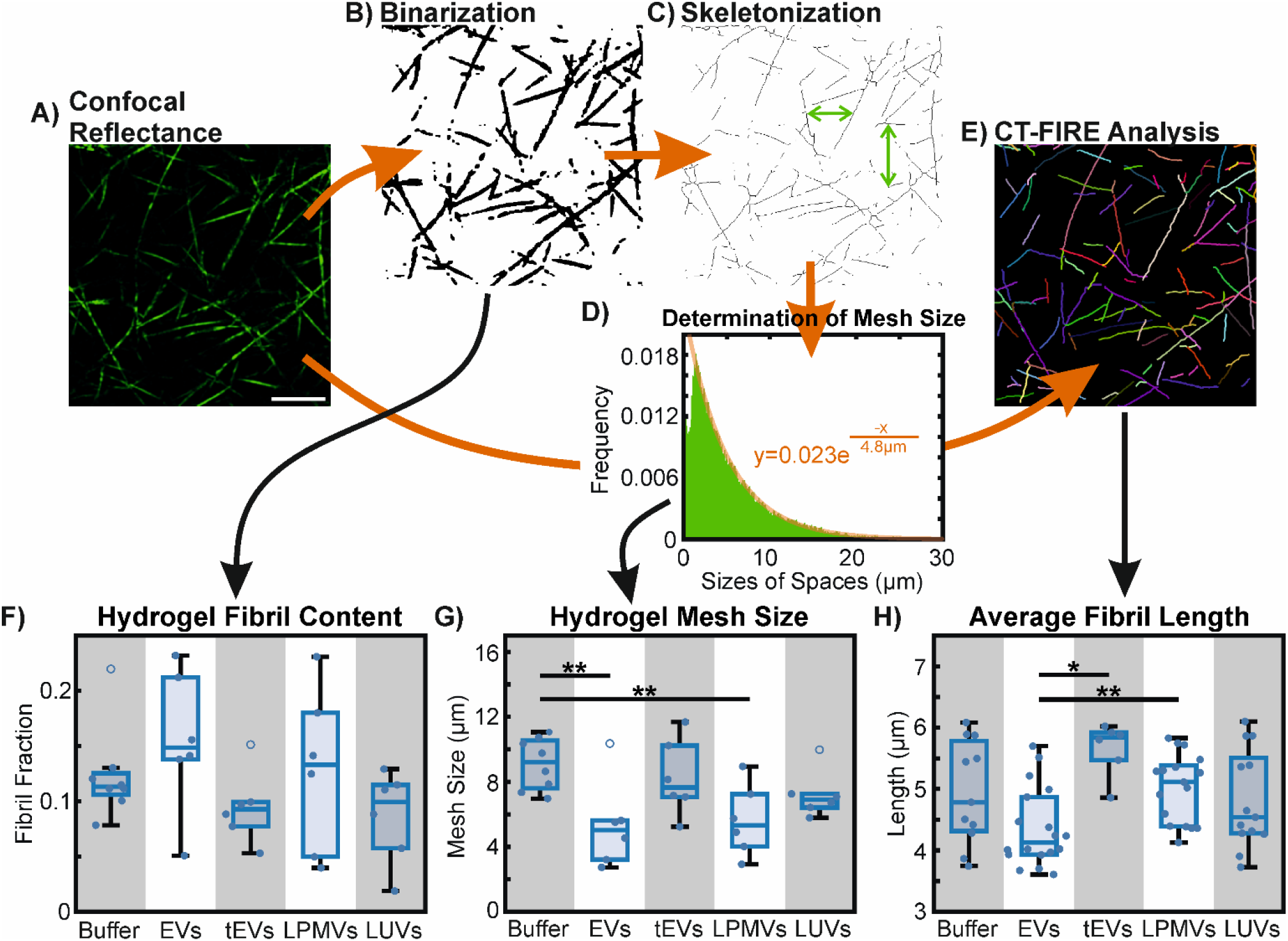
Analysis of collagen I fibril geometry and hydrogel mesh structure. A) Confocal reflectance microscopy is used to image collagen I fibrils in hydrogels. In this example, collagen fibrils are formed in calcium-containing buffer (HBS+Ca) without any particle inclusions (Buffer control). Scale bar: 10 µm. B) Images are binarized and the proportion of ‘fibril pixels’ is summed up as a fraction of total pixels to obtain hydrogel fibril content. C) Binarized images are further skeletonized to obtain the central axes of the fibrils. D) Spaces between the fibrils are counted up in the xand y-directions and displayed in histogram form, where they fall in an exponential distribution. Distributions are fitted to exponential curves to determine their mean values, which are converted to hydrogel mesh size in µm. E) CT-FIRE analysis (48) is used to identify and determine the length of collagen fibrils (shown as overlays on the original image). F) Fibril content of hydrogels formed with EVs, tEVs, LPMVs, and LUVs, compared to particle-free gels. No statistically significant differences were found with 1-way ANOVA (p>0.05) with n=8 replicates for the particle-free control (Buffer) and n=6 replicates for other conditions. G) Hydrogel mesh size, i.e. the average amount of space between fibrils. Statistically significant differences were determined with 1-way ANOVA and Tukey-Kramer post-hoc analysis over n=8 replicates for the particle-free control (Buffer) and n=6 for other conditions, as indicated by ** (p<0.05). H) Average length of fibrils identified in gels containing different particles. Statistically significant differences were determined with 1-way ANOVA and Tukey-Kramer post-hoc analysis over n=11, 18, 6, 17, and 13 replicates for Buffer, EVs, tEVs, LPMVs, and LUVs, respectively. Significance levels are indicated by * (p<0.01) and ** (p<0.05). Individual points represent independent experimental replicates.

We found that differences in hydrogel rheology in the presence of calcium and EVs are reflected in fibril geometry and matrix structure. Overall fibril content of hydrogels is not significantly affected by the presence of any particle, although EVs and LPMVs appear to slightly increase fibril content. Hydrogel mesh size is significantly decreased by EVs and LPMVs, but not by tEVs or LUVs when compared to particle-free gels. Finally, average fibril length does not appear to be affected significantly by particles when compared to particle-free gels, but the presence of EVs appears to result in significantly shorter fibrils compared to gels with tEVs and LPMVs. It is possible that this shorter fibril length in EV-containing gels is in part due to greater intersection of fibrils, causing the CT-FIRE software to detect multiple shorter fibril sections as opposed to single continuous fibrils. Qualitatively, upon visual inspection, hydrogels formed with EVs appear to have both shorter fibrils and more intersecting fibrils compared to other conditions. With unchanged, or even slightly increased fibril content combined with smaller mesh size and shorter average fibril length, it follows that gels formed with EVs would have a greater absolute number of fibrils (or fibril sections) per unit volume. Fibril number could not be accurately estimated with CT-FIRE analysis due to inconsistencies in fibril segmentation.

Taken together with our rheological data, our analyses of hydrogel and fibril structure suggest that EVs accelerate the gelation process by nucleating fibrils and promoting network formation. The shortened nucleation phase duration and the slight increase in fibril content of formed gels is indicative of EVs helping to nucleate new fibrils. The increase in peak G’ growth rate and greater final G’ value suggests that EVs also promote and accelerate network formation. This appears to be reflected in the smaller mesh size and shorter detected fibril length, which suggest more nucleated fibrils and a greater degree of intersecting and overlapping fibrils, resulting in a more densely packed fibril matrix. LPMVs also decrease the nucleation phase duration and decrease the hydrogel mesh size, suggesting a similar ability to nucleate fibrils compared to EVs. Because LPMVs do not increase the G’ growth rate as much as EVs, they likely are not as effective in promoting network formation. This is reflected in the longer average detected fibril length compared to gels formed with EVs. This suggests that EVs, with their distinct membrane composition, may have a specialized function in collagen I fibrillogenesis that cannot be achieved with whole cell plasma membrane.

In contrast to native EVs, trypsinized tEVs behave much more like synthetic LUVs, in that they do not stimulate the gelation process or affect the resulting fibrils and matrix structure. This suggests that simply having a particle inclusion during the gelation process does not significantly aid in collagen I fibril nucleation and that functional proteins are largely responsible for interactions between EVs (and to some extent, LPMVs) and collagen I. Compared to EVs, collagen gels formed with tEVs have significantly longer fibrils and are not significantly different from gels containing LPMVs, LUVs, or particle-free controls. One possible reason why the fibrils are so long could be due to non-specific electrostatic interactions. The overall surface charge of EVs (measured as zeta potential; Suppl. Fig. S3) does not change upon trypsinization, but the trypsin enzyme is known to target charged amino acid residues (49), which may result in ragged charged protein fragments. In the presence of a divalent cation like calcium, such charged fragments may act to ‘bridge’ collagen fibrils, resulting in longer fibrils overall.

### EVs become integrated into the collagen I fibril matrix

Further analysis of confocal images was conducted to investigate particle localization in collagen hydrogels with respect to their fibril matrix structure. Specifically, we quantified the particleto-fibril distance and the proportion of fibril-associated particles, as done previously (32). Fluorescently labelled EVs, tEVs, LPMVs, and LUVs were imaged alongside the confocal reflectance microscopy of collagen I fibrils (Fig. 4A). By binarizing the images of the particles and determining their geometric centres of mass (Fig. 4B), we were able to cross-reference their locations with the central axes of the collagen fibrils using the Python implementation of the KD-Tree nearest neighbour search algorithm (50). This enabled us to determine the minimum distance from each particle centre to its nearest fibril. We also defined a fibril-associated particle to be a particle whose centre of mass falls within 500 nm of the central axis of a collagen fibril (Fig. 4C). This is approximately the noise floor for determining particle-fibril colocalization, being half the apparent width of a fibril (2-3 pixels or 200-300 nm) and half the apparent radius of an average particle as they appear in our images (also 2-3 pixels, or 200-300 nm). We found that on both metrics, EVs behaved differently compared to tEVs, LPMVs, and LUVs. On average, EVs localize significantly closer to collagen fibrils than tEVs and LUVs and have a significantly greater proportion of fibril-associated particles. This is partly due to there being more densely-packed fibrils, and thus a smaller average mesh size in collagen gels formed in the presence of EVs compared to tEVs and LUVs (Fig. 3F,G). It is also evident that tEVs behave similarly to LUVs with relatively low levels of colocalization by the two metrics analyzed, further supporting our hypothesis that proteins are responsible for EV-collagen matrix interactions.

**Figure 4.**
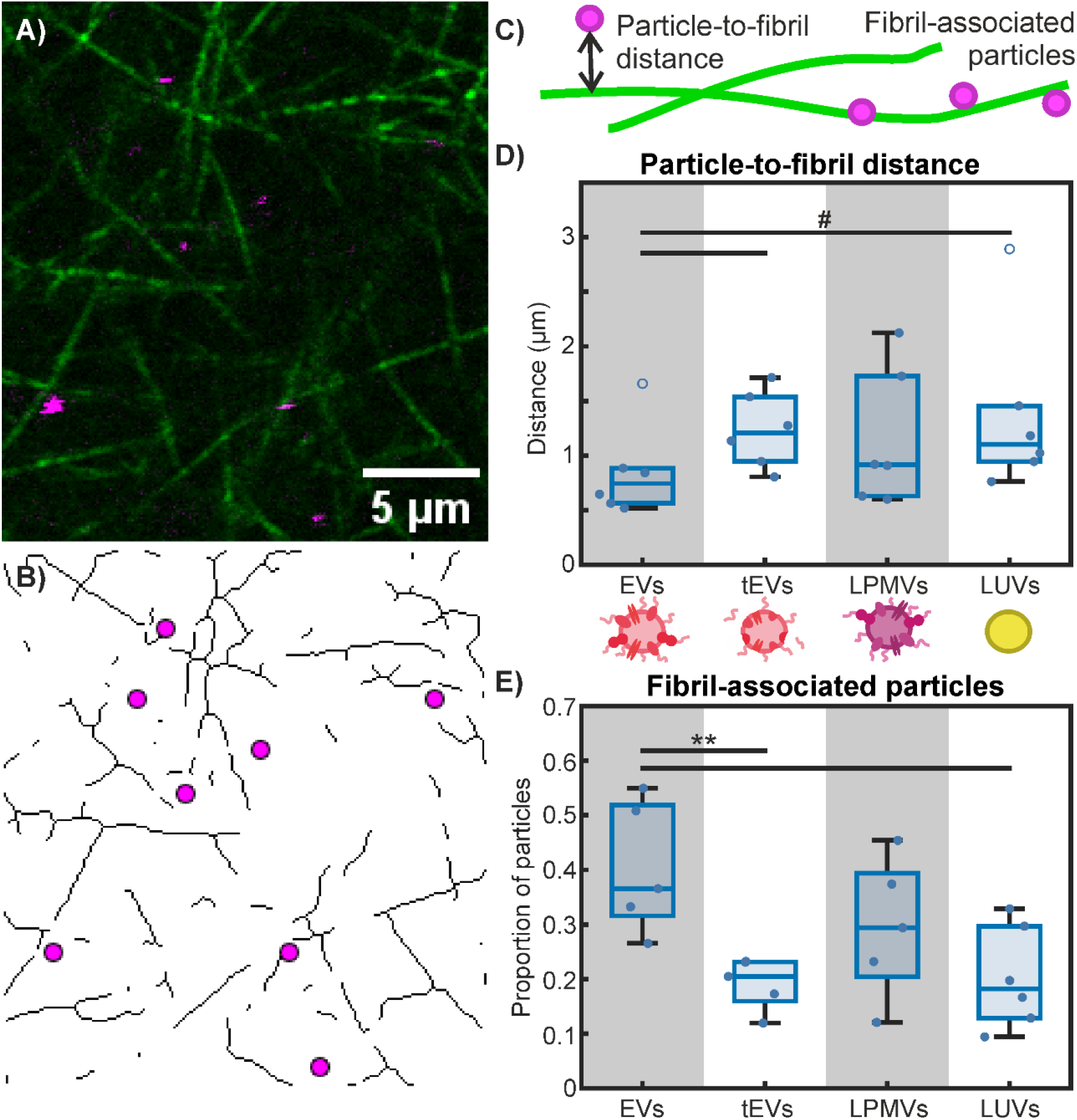
Particle localization within collagen I hydrogels. A) Representative image of collagen I fibrils (green) imaged with confocal reflectance microscopy and fluorescently labelled EVs (magenta), imaged with fluorescence confocal microscopy. B) The image from (A) is binarized and skeletonized to obtain the central axes of the collagen fibrils. The geometric centres of mass of the particles are determined and shown as magenta circles. C) Schematic diagram showing particle-to-fibril distance and fibril-associated particles. The former refers to the minimum distance between a particle’s centre of mass to the nearest fibril central axis and the latter refers to any particle whose centre of mass lies within 500 nm of a fibril’s central axis. D) Average particle-to-fibril distance of EVs, tEVs, LPMVs, and LUVs. A 1-way ANOVA test revealed no statistically significant differences across conditions, but 2-way t-tests were conducted on isolated data between EVs and tEVs, and EVs and LUVs to see if any differences could be found. Significant differences (p<0.01) found this way are indicated with #. E) Proportion of particles determined to be associated with collagen fibrils. Outlier data points that were excluded from statistical analyses are indicated with empty blue circles. Statistically significant differences were determined with 1-way ANOVA with Tukey-Kramer post-hoc analysis, as indicated with ** (p<0.05).

Despite having similar, albeit slightly reduced effects on gelation kinetics and matrix structure, LPMVs do not appear to associate as strongly or persistently with collagen I fibrils as EVs do. This suggests weaker or possibly more transient interactions, whereby LPMVs could help to nucleate fibrils and then detach. These functional differences, again, serve to emphasize the specialized composition and role of EVs as structures distinct from the plasma membrane of their source cells.

EVs are known to become entangled and entrapped in ECM structures, where they provide important contextual cues to resident cells (51– 53). The question of how EVs come to be so intimately integrated into ECM can be answered by two possible scenarios: EVs could migrate into fully-formed ECM environments whereupon they become entrapped, or new ECM can be built up around EVs, trapping them in place. The latter possibility implies a mechanism by which EVs can actively partake in ECM formation and remodeling, which our data thus far supports. To further investigate this aspect, we compared the colocalization of EVs embedded in collagen I hydrogels by having them present prior to gelation with that of EVs infiltrating and diffusing into pre-formed hydrogels (Fig 5A,B). We previously reported on EVs infiltrating into pre-formed collagen I hydrogels (32), and re-use some of that data here for our present analysis.

**Figure 5.**
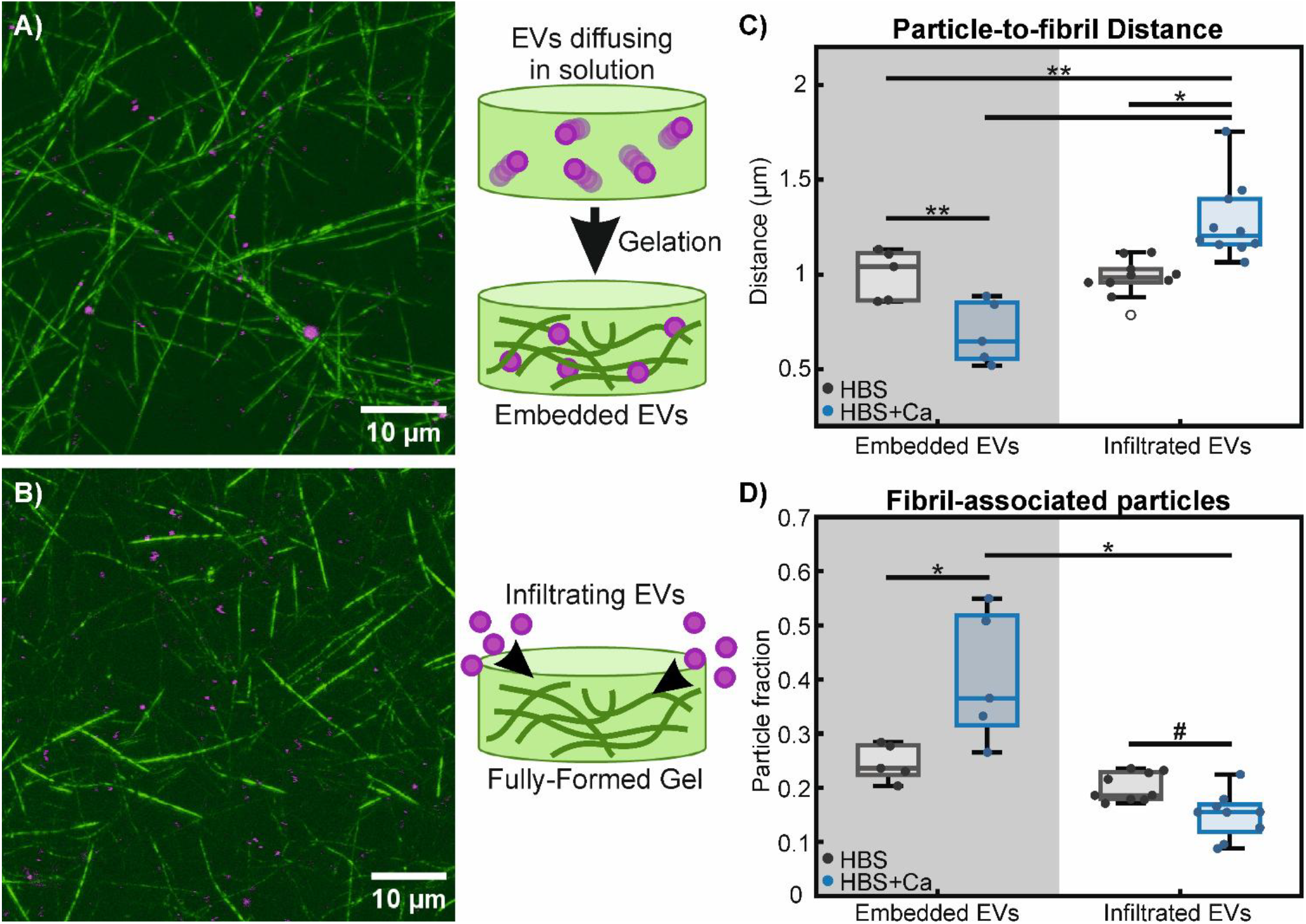
EVs take part in fibrillogenesis and become integrated into the matrix structure. A,B) Combined confocal reflectance microscopy of collagen I fibrils (green) and confocal fluorescence microscopy of labelled EVs (magenta). Images are vertical projections of 10-image stacks (obtained from larger data sets containing 30 total images), covering a total vertical depth of 7 μm in order to better visualize the fibril matrix structure. In A, EVs were embedded into the hydrogel by mixing them with the collagen solution prior to gelation, as shown in the schematic. In B, EVs are allowed to diffuse and infiltrate into fully-formed collagen gels, as shown on the right. Imaging data for EVs infiltrating into pre-formed gels (including B) are reused from a previous report on EV diffusion (32). C) Average particle-to-fibril distance of embedded and infiltrated EVs, in both calcium-free (HBS) and calcium-containing (HBS+Ca) buffers. D) Proportion of fibril-associated particles for embedded and infiltrated EVs in calcium-free and calcium-containing buffers. A 2-way ANOVA test determined significant differences (p<0.05) between embedded and infiltrated particles, but not due the presence of calcium in the buffer. Further testing with 1-way ANOVA and Tukey-Kramer post-hoc analysis found significant pairwise differences, as indicated with * (p<0.01) and ** (p<0.05). Significant differences determined by 2-way t-test on isolated data are indicated with # (p<0.01). Outlier data points that have been excluded from statistical analyses are indicated with empty circles. Experiments consist of n=6 replicates for embedded EVs and n=10 for infiltrated EVs.

Qualitatively, upon visual inspection of images, embedded EVs appear to colocalize well with fibrils and seem to be associated with points of intersection between fibrils or with adjacent parallelly-aligned fibrils (Fig. 5A). This does not seem to be the case with infiltrated EVs, which appear randomly distributed among fibrils and mesh spaces. This is corroborated with quantitative measurements of particle-to-fibril distance and proportions of fibril-associated particles (Fig. 5C,D). Embedded EVs have a lower average particle-to-fibril distance and a higher proportion of fibril-associated particles than infiltrated EVs, but only in the presence of calcium. Without calcium present, no significant difference is observed. Our data suggest that EVs do not interact with fibrillar collagen, at least not with the same strength as with colloidal collagen I in solution. EVs simply diffusing into fully-formed collagen hydrogels are not able to integrate into the fibril matrix. Moreover, when calcium is present, infiltrated EVs appear to have even lower colocalization with fibrils, possibly due to having increased affinity for non-fibrillar, unincorporated collagen molecules in the mesh spaces (54). In contrast, our imaging and rheological data show that embedded EVs within solutions of colloidal collagen I and in the presence of calcium actively engage with collagen molecules, aiding in their nucleation and growth into fibrils, becoming intimately integrated into the resulting fibril matrix structure.

### Possible underpinnings of EV-collagen interactions

Our previous work has shown that argynylglycylaspartic acid (RGD) motif-binding integrins contribute to the immobilization of EVs in collagen I hydrogels (32), and thus their overall mobility in such an environment. Importantly, RGD motifs in the collagen I molecule are only available for binding in a partially or fully-denatured, non-fibrillar state (55, 56). A similar interaction appears to be involved in our present investigation. The calcium dependence of the phenomena that we have investigated strongly suggest the involvement of RGD-binding integrins, as cations – and calcium, specifically, are required for the dimerization of functional integrin complexes that bind the RGD motif (56, 57). What remains unclear is whether these initial integrin-RGD interactions persist after fibril formation, or if other molecules are involved in the continued close association of EVs with collagen I fibrils that we observe. It is possible that EVs possess other integrin complexes that can bind to fibrillar collagen once they are brought into close enough proximity, such as those that bind the GFOGER motif (58–60). Such complexes may require divalent cations other than calcium, however, to properly dimerize (56, 57). This specificity for certain cations could potentially be used to narrow down what specific integrin complexes are present in EV membranes (59, 61). Indeed, it may even be possible to tune relative EV affinities for fibrillar versus non-fibrillar collagen or other ECM materials with different ions in the medium, as seems to be the case for infil-trated EVs (Fig. 5D).

### Implications of EV-mediated fibrillogenesis

Although our findings represent in vitro phenomena in minimal experimental conditions, our data nevertheless have important implications in real physiological and pathological processes. Systematic study of interactions between EVs and purified ECM components can lead to a better understanding of how these interactions contribute to EV interactions with whole tissue.

Mounting evidence shows that EVs, whether directly or indirectly, are involved in the formation and dynamic evolution of ECM. Moreover, our data provide a plausible mechanism by which EVs themselves become an integral component of the tissue microenvironment. By becoming embedded into the matrix architecture, EVs become an important signaling cue for resident cells (51–53). Indeed, this may be a crucial aspect in physiological, as well as pathological contexts. In wound healing, for example, EVs from various cell sources carry important molecular payloads that help promote and direct angiogenesis, cell proliferation, and ECM remodeling processes (6, 62– 64). Coordination of such a complex biological process involving many different cell types and matrix molecules requires spatiotemporal control of signaling processes (64, 65). From the initial production of provisional matrix to stabilize the wound, to its gradual replacement with granulation tissue, and finally with the restoration of normal tissue – the ECM in the healing wound is in dynamic flux (66, 67). Matrix-bound EVs would, thus, be deposited and embedded into the microenvironment, only to be released to deliver their payloads according to the active remodeling processes taking place. This ‘controlled release’ aspect of EV-matrix association is potentially exploitable in tissue engineering and regenerative medicine applications (51, 52, 68, 69). EVs may also play a role in the initial stages of wound repair, as the ability to recruit and incorporate partiallyto fully-denatured collagen molecules generated during injury into provisional ECM structures may help to stabilize wounds in preparation for further remodeling and healing processes. To this end, our results showing the acceleration of fibrillogenesis in a collagen hydrogel and the integration of EVs into the matrix structure could contribute to the development of new bioactive scaffolds or of wound-stabilizing, collagen-crosslinking EV suspensions.

Our data showing that breast cancer cell-derived EVs – and specifically those of an aggressive, metastatic, triple-negative cell phenotype, are capable of nucleating and aiding in the formation of new collagen I fibril structures also has interesting implications for cancer pathologies. EVs promoting and accelerating fibrillogenesis may contribute to fibrotic reorganization of the cancer niche. These effects might further translate to overall tissue stiffening and fibrosis *in vivo*, often associated with solid tumours (70–72). Dysregulation of fibrillar collagen networks contribute to Hanahan and Weinberg’s ‘Hallmarks of Cancer’ (73, 74), cellular characteristics and processes that define and enable cancerous growth. This is best appreciated at the tissue level, where changes in tissue collagen content are implicated in a wide range of effects, from modulating immune reactivity (75, 76), to enabling the migratory and invasive behaviour of metastatic cells (72, 77–79), or rewiring signaling pathways that govern cell behaviour and survival (30, 80–83). This is on top of the continually gro-ing body of evidence of EVs directly reprogramming cells to promote cancer cell malignancy (3, 5, 38, 84, 85).

## Conclusions

Our investigation into the interactions between breast cancer cell-derived EVs and soluble collagen I has shown that EVs can directly participate in the formation of new ECM structures. Rheological measurements show that the gelation process of collagen hydrogels is affected during both the initial nucleation phase, as well as during the growth and extension of nucleated aggregates into fibrils. We show that the calcium-dependent interactions between EVs and collagen are mediated by membrane proteins that are ablated by treatment with trypsin. Image analysis of collagen gels shows that EVs nucleate matrices composed of shorter, more densely packed fibrils. In taking part in collagen I fibrillogenesis, EVs become integrated into the resulting fibril matrix. Analyses of particle-fibril colocalization show that this close association of particles with fibrils cannot be achieved by simply having EVs diffusing into fully-formed hydrogels, but only arises by the active participation of EVs in collagen fibrillogenesis.

Our work shows that EVs are not merely passive messengers of cell signals, but themselves play an active role in the dynamic molecular processes involved in ECM formation and remodeling. This has wide implications in tissue engineering and regenerative therapy applications, as well as in cancer research. As bioactive components of tissue microenvironments, EVs can potentially be exploited, not only as mediators of cell signaling, but also as integral components of new bioactive scaffolds that can aid in matrix synthesis and regeneration. EVs evidently have greater clinical value than merely being diagnostic or prognostic indicators. In cancer, their ability to alter the local tumour microenvironment, as well as distant tissue sites means EVs enable disease progression and the development of metastatic lesions. This aspect of cancer biology may need to be addressed by future holistic anti-cancer therapies.

Our key findings are that (i) EVs accelerate collagen I fibrillogenesis and help to form more densely packed fibril matrices with greater final storage and loss moduli than particle-free controls; (ii) intact proteins are required for the calcium-dependent interactions between EVs and collagen I; (iii) EVs become integrated into the matrix and are closely associated with fibrils; and

(iv) this association of EVs with fibrils cannot be achieved by simply having EVs diffusing into a pre-formed collagen I gel.

## Materials and Methods

### Cell culture

MDA-MB-231 breast cancer cells were obtained from the American Type Culture Collection and cultured in 10 cm-diameter tissue culture polystyrene Petri dishes (Nunc™ Cell Culture Treated Plates; Thermo Fisher Scientific, USA) at 37 °C under 5% CO2. The complete culture medium consisted of low-glucose Dulbecco’s Modified Eagle’s Medium (DMEM; Sigma-Aldrich, USA) supplemented with 10% fetal bovine serum (FBS; Thermo Fisher Scientific, USA) and 1% penicillin-streptomycin (Thermo Fisher Scientific, USA). Cells were passaged every 3-4 days at 80-90% confluency as follows: old medium was removed and the plates rinsed twice with phosphate buffered saline (PBS; 137 mM NaCl, 2.7 mM KCl, 10 mM Na_2_PO_4_, 1.8 mM KH_2_PO_4_). Cells were detached from the plates by incubating with 2 mL trypsin/EDTA solution (PAN-Biotech, Germany) for 5 minutes at 37 °C. The trypsin was quenched with 2 mL complete culture medium and the cell suspension was collected and centrifuged at 200 rcf for 10 minutes. The supernatant was discarded and the pelleted cells were re-suspended with fresh medium. Cells were plated on Nunc cell culture-treated Petri dishes (Thermo Fisher Scientific, USA) with 7-8 mL of complete culture medium at an approximate split ratio of 1 to 3 or 4.

### Buffers

Two different buffers were used to study the calcium-dependence of EV-collagen interactions. It was not possible to use PBS, as the addition of calcium caused precipitation of insoluble calcium phosphate. Instead, a HEPES-based buffer was used, according to a previously-published protocol for generating giant plasma membrane vesicles (GPMVs) (46). In the original publication, this buffer was referred to as GPMV buffer and consists of 150 mM NaCl, 10 mM HEPES, and 2 mM CaCl2; pH 7.4. Here, we refer to this buffer as HBS+Ca to contrast it from our calcium-free buffer. The concentration of calcium used reflects nominal extracellular calcium levels *in vivo* (86, 87). Our custom calcium-free HEPES-buffered saline (HBS) was developed to match the osmolality of HBS+Ca, substituting the CaCl_2_ with more NaCl: 150 mM NaCl + 16 mM NaCl, pH 7.4. The osmolality of both buffers was determined to be 303 mOsm/kg using a Gonotech feezing point osmometer (Gonotech, Germany).

### Generation and purification of EVs

To obtain enough EVs for experiments, 10-12 plates (10 cm-diameter) of cells were cultured to 80-90% confluency. Cells were rinsed 3 times with PBS before the medium was replaced with 8 mL serum-free medium per plate, consisting of low-glucose DMEM + 1% penicillin/streptomycin. Removing FBS from the medium prevented contamination with bovine vesicles and induced a state of serum starvation, which should have enhanced EV production (88). The medium was conditioned by incubating it with cells over 3 days. The conditioned medium was then collected and centrifuged at 400 rcf for 10 minutes to pellet dead cells. The pellet was discarded and the supernatant centrifuged again at 2000 rcf to remove remaining cell debris. This supernatant was then concentrated using 100 kDa Amicon Ultra-15 centrifugal filters (Merck Millipore, USA), centrifuged at 3400 rcf to a final volume of 1 mL. The concentrated conditioned medium was then incubated for 10 minutes with 1 µL 2.5 mg/mL 1,1’-Dilinoleyl3,3,3’,3’-Tetramethylindocarbocyanine, 4-Chlorobenzenesulfonate (FAST DiI; Fischer Scientific, USA) dissolved in ethanol at 37 °C to label the lipid structures. EVs were then separated with SEC using Sepharose CL-4B base matrix (Sigma-Aldrich, USA), as previously reported (32, 34, 36) and described below.

To equilibrate the Sepharose, 15 mL of suspended Sepharose matrix was allowed to settle over 2 hours at 7 °C and the liquid medium was decanted off and replaced with HBS. This was repeated 5 times to wash the Sepharose beads. A column was prepared using a 10 mL syringe with the plunger removed. The end was stopped with an end cap and a Whatman polycarbonate membrane filter of 10 µm pore size (Sigma-Aldrich, USA) was cut and placed in the bottom of the syringe to prevent the Sepharose from running out. After the final wash, Sepharose beads were suspended in HBS, loaded into the prepared syringe, and left overnight to pack. Two column volumes (20 mL) of HBS were run through the column before the sample was loaded. Up to 20 fractions of approximately 500 µL each were collected and eluted with additions of 1 mL HBS at a time. In a previous report, we showed that fractions 7-10 are enriched in 100-400 nm-diameter particles. Dynamic light scattering (DLS) data was included in the supplemental materials of our previous paper (32) and have been included in Suppl. Fig. S1. These fractions were pooled together and re-concentrated using centrifugal filter tubes with a molecular weight cutoff of 100 kDa. To obtain particles in HBS+Ca, suspensions were concentrated with centrifugal filters to 200 µL, then resuspended in HBS+Ca to 800 µL. This was repeated 5 times to gradually wash out the old buffer. EV suspensions were aliquoted, frozen, and stored at −80 ^°^C with 10% dimethylsulfoxide (DMSO; Sigma-Aldrich, USA) as a cryopreservant in HBS and were thawed prior to use in experiments.

To determine relative EV concentrations (and also for other particles), a 10 μL droplet of particle suspension was imaged using a pco.Edge sCMOS camera (PCO AG, Germany) mounted to a Zeiss AXIO Observer.D1 microscope equipped with a 63× 1.2 NA water immersion C-Apochromat objective (Carl Zeiss, Germany) in epifluorescence mode with the built in filter set for Texas Red fluorescence. Particles were identified and counted in FIJI. Briefly, images were thresholded and binarized to include only pixels corresponding to particles. Particles were then counted with the built-in ‘Analyze Particles’ function. Absolute concentrations in units of particles per volume are difficult to obtain due to the nature of the microscopy used and lack of depth information. Regardless, this pseudo-2D particle per unit area count allowed dilution of suspensions to maintain approximately the same particle concentration in experiments. The particle concentration in units of particles per volume can be estimated by dividing the pseudo-2D concentration by the depth of field of our imaging setup. This was experimentally determined to be ~700 nm by observing the range in which the fluorescence signal from particles adhered to the surface of a microscope slide was acceptably above the background noise while varying the vertical focus knob both above and below the plane of the particles. This agrees well with an estimate based on a theoretical calculation (89). The final in-gel concentration of particles corresponding to the relative concentration of 1 in Fig. 1 and also to the final in-gel concentration of particles used for all other experiments was thus 0.026±0.004 particles×μm^-3^ or 26±4 million particles×μL^-1^.

### Trypsinization of EVs

To determine whether EV interactions with collagen I were due to membrane proteins, EVs were treated with trypsin to digest their surface proteins (tEVs). EV suspensions corresponding to EVs from 10-12 plates’ worth of cells incubated over 3 days in serum-free medium were first concentrated from 1mL to 100 μL with centrifugal filter tubes with a molecular weight cutoff of 100 kDa. Next, 200 μL of TrypLE™ Express Enzyme (Thermo Fisher, USA) was added directly to the suspension in the filter tube, which was then incubated for 10 minutes at 37^°^C. This was then quenched by diluting the solution with HBS (or HBS+Ca) to fill the tube (approximately 800 μL). The tube was centrifuged at 3400 rcf to wash out the excess buffer and trypsin, concentrating the suspension down to 100 μL. The tube was refilled with buffer and washed in this way a further 4 times to remove the trypsin and replace the buffer. Particle concentration after trypsinization was checked and adjusted as described for EVs just prior to experiments.

### Generation of GPMVs and extrusion of LPMVs

LPMVs were produced, as described previously (32). GPMVs were first produced as follows (46): 10-12 plates of 80-90% confluent cells were washed twice with PBS, then once with HBS+Ca. Vesiculating medium was made by adding 2 µL of a 1 M N-ethylmaleimide (NEM; Sigma-Aldrich, USA) stock solution dissolved in water per 1 mL of HBS+Ca. Each plate of cells received 2 mL of vesiculating medium and were left to incubate at 37 °C for 1 hour to allow for GPMV formation. GPMVs were then collected by gentle pipetting to avoid lifting up the cells. This suspension was centrifuged at 100 ×g to pellet cell debris. The pellet was discarded and the supernatant centrifuged for 1 hour to pellet the GPMVs. This supernatant was discarded and the GPMV pellet resuspended in 1 mL HBS+Ca. The GPMVs were then incubated with 1 µL FAST-DiI for 10 minutes at 37 °C to stain them. Labelled membranes were next extruded using an Avanti handheld extruder fitted on a heating block (Avanti Polar Lipids, USA) set on a hot plate at 37 °C, first 21 passes through a Whatman Nuclepore 400 nm-pore size track-etched membrane filter, then 21 passes through a 200 nm-pore size filter (Sigma-Aldrich, USA). The resulting LPMVs were concentrated and (when needed) the buffer changed as with EVs using centrifugal filters. Particle concentrations were checked and adjusted as described for EVs.

### Production of LUVs

LUVs were produced from lipid mixtures consisting of 4 mM DOPC (Avanti Polar Lipids, USA) dissolved in chloroform with 0.5 mol% DiI (Fisher Scientific, USA). First, 30 µL of lipid was spread in a clean glass vial and dried under vacuum for 1 hour. Next, the lipid layer was rehydrated in 1 mL HBS or HBS+Ca, then vortexed for 5 minutes to form multilamellar membrane structures. The resulting suspension was then extruded 21 passes with a handheld extruder fitted with a 200 nm-pore size Whatman Nuclepore track-etched membrane filter. Particle concentrations were checked and adjusted as described for EVs.

### DLS analysis of size and zeta potential

DLS was used to ensure consistency in size and charge of EVs, LPMVs, and LUVs. Particle suspensions were loaded into disposable folded capillary tubes (DTS1070; Malvern Panalytical, UK) and analyzed with a Malvern Instruments Nano-ZS Zetasizer equipped with a 632.8 nm 4 mW HeNe laser (Malvern Panalytical, UK). Size measurements were obtained with a scattering angle of 173^°^ prior to determination of zeta potential. Absolute measurement of surface charge was not possible due to electrostatic screening from the high salt content of the buffers used. Thus, the measured zeta potential was used to illustrate relative surface charge between particle types.

### Rheology of collagen I gelation

Bulk rheology was conducted on an Anton Paar MCR301 rheometer with a 12 mm cone-plate (CP12) geometry probe (Anton Paar, Austria) in oscillatory mode. Collagen I solutions were mixed directly on the rheometer stage, cooled to 7 ^°^C to prevent premature gelation and with the reservoir of the stage filled with distilled water to maintain humidity. First, 12.5 μL of a 6 mg/mL stock solution of solubilized collagen I from bovine skin (Sigma-Aldrich, USA) was pipetted onto the stage, followed by 1 μL 1 M NaOH to bring the pH up to approximately 7. This was then diluted 1:1 with 12.5 μL 2× concentrated buffer (HBS or HBS+Ca) and mixed by pipetting up and down. This was further diluted 1:1 with 24 μL 1× buffer, resulting in a final collagen concentration of 1.5 mg/mL. The sample volume was reduced by discarding 30 μL of this mixture in order to better fit the rheometer probe. Solutions were mixed in this manner instead of with smaller volumes due to the viscosity of the collagen making pipetting of smaller volumes unfeasible. The probe was lowered onto the liquid sample to its manufacturer-determined measurement position before the stage was heated to 35 ^°^C and the Peltier hood lowered on top. A time sweep was then immediately started to measure the storage and loss moduli (G’ and G”, respectively) at 30 s intervals with 1% strain and 1 Hz oscillation for 40 minutes. The sample reached the target temperature of 35^°^C by the second measurement point (approximately 1 minute). Once the time sweep was complete, the collagen mixture was allowed to sit an extra 20 minutes to ensure complete gelation before an endpoint frequency sweep from 0.1 to 10 Hz with 1% oscillatory strain was conducted to determine the final storage and loss moduli.

Analysis of rheology data was conducted using MATLAB. The endpoint storage and loss moduli were determined by averaging the values over 0.16 to 2.5 Hz, which was the frequency range over which G’ and G” remained relatively constant. For the time sweep gelation kinetics data, the first derivative of the storage modulus data was determined and plotted, as shown in Fig. 1. We defined the end of the nucleation phase as follows: for each individual gelation curve, the first 5 time points of 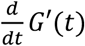 were used as a baseline to determine the mean and standard deviation. The end of the nucleation phase was then the time point at which the value of the first derivative exceeded 2 standard deviations of the baseline mean value for two consecutive time points. The peak growth rate of the storage modulus was the maximum point of the first derivative over time, which coincides with the inflection point of the storage modulus.

To determine lack of statistically significant dose-dependence of rheological parameters, the Pearson’s correlation coefficient was calculated between normalized particle concentrations and nucleation phase duration, as well as peak G’ growth rate using MATLAB.

### Formation and confocal imaging of collagen I gels

For confocal imaging, collagen gels were mixed and formed in 96-well plates with glass bottoms of 0.2 mm thickness (Greiner Bio-One, Austria). Plates and reagents were kept on ice to prevent premature gelation of the collagen. First, 12.5 μL of 6 mg/mL soluble collagen I stock was added to each well, followed by 1 μL 1 M NaOH to bring the pH up to approximately 7. To this, 12.5 μL 2× buffer (HBS or HBS+Ca) was added and mixed by pipetting up and down. Next, 2 μL of particle suspension was added. This was further diluted with 22 μL 1× buffer to obtain a final volume of 50 μL and final collagen I concentration of 1.5 mg/mL. Once mixed, the plate was transferred to an incubator and allowed to gel at 37 ^°^C overnight.

For experiments involving infiltrated particles, gels were formed without the addition of particles, replacing them with an equal amount of 1× buffer. This was allowed to gel overnight at 37 ^°^C before 10 μL of particle suspension was pipetted directly on top of each gel and allowed to diffuse throughout for a minimum of 3 hours at 37 ^°^C.

The fibril structure of collagen hydrogels was imaged in confocal reflectance mode with a Leica SP8 microscope equipped with a 63× 1.2 NA water immersion objective (Leica, Germany) with 488 nm argon laser illumination. Z-stacks consisting of 30 images were obtained with 0.75 μm spacing to obtain more data than could be obtained from a single slice image. For gels containing DiI-labelled particles, particles were imaged in parallel with excitation from a 561 nm wavelength diode laser.

Image analysis of fibril content, mesh size, and fibril length is illustrated in Fig. 3. Images were first processed with a bandpass filter to remove scanner artefacts. For fibril content and mesh size, images were thresholded and binarized in FIJI. Fibril content was determined by summing the fibril pixels and dividing by the total number of pixels in the image. To determine average mesh size, or the average amount of space between collagen fibrils, we used a previously described process (41, 47) with slight modification: the binarized images were first skeletonized in FIJI. Images were then processed in MATLAB to determine the number and size of spaces between fibrils in the x- and y-directions, which, when plotted in a histogram of sizes of spaces, fall into an exponential distribution. The mean mesh size value corresponded to the value of the exponential coefficient of a fitted exponential function, converted from pixels to micrometers. For the determination of fibril length, we processed unbinarized images with the CT-FIRE collagen fibril analysis software developed by Eliceiri et al. (48).

For the determination of particle-to-fibril distance and the proportion of fibril-associated particles, images of DiI-labelled particles were binarized in FIJI and the coordinates of the geometric centres of mass of particles were determined with the ‘Analyze Particles’ function. These coordinates were cross-referenced with the positions of binarized, skeletonized collagen fibrils and the distance from each particle centre to its nearest fibril was determined using the Python implementation of the KD-Tree nearest neighbour search algorithm (50). Analysis was done in pseudo-3D, whereby adjacent slices in the 3D stack were also considered in the search for the nearest neighbouring fibril, but not any further due to the stack step size being sufficiently large that fibrils more than 1 slice away could be ignored. Fibril-associated particles were defined to be particles whose centres of mass lie within 500 nm of the central axis of a collagen fibril, corresponding approximately to the noise floor of particle-fibril colocalization: the apparent average radius of particles, as they appeared in images (2-3 pixels or 200-300 nm) plus half the apparent radius of the collagen fibrils (also 2-3 pixels or 200-300 nm).

### Statistics

Unless otherwise specified, individual experimental replicates consist of independently formed collagen gels, with embedded particles deriving from multiple rounds of EV collection and treatment or LUV formation to ensure reproducibility. Statistical tests were carried out using MATLAB software, including 2-way t-tests and 1-way and 2-way ANOVA with Tukey-Kramer post-hoc analyses, as indicated.

## Acknowledgments

N.W. Tam would like to acknowledge funding from the International Max Planck Research School on Multi-Scale Biosystems. A. Cipitria would like to acknowledge funding from the DFG Emmy Noether grant (CI 203/2-1), from IKERBASQUE Basque Foundation for Science, from Fundación Científica Asociación Española Contra el Cáncer (grant LABAE223466CIPI), from the Spanish Ministry of Science and Innovation (MCIN/AEI/10.13039/501100011033/FEDER UE, through grant PID2021-123013OB-I00) and from the European Research Council Consolidator Grant (DORMATRIX, 101123883).

## Data Availability

All necessary raw and processed data for reproducing our findings, as well as MATLAB and Python scripts used for our analyses can be found in the publicly accessible Edmond repository of the Max Planck Society (https://doi.org/10.17617/3.AMI3GV).

## Conflict of Interest Disclosure

The authors declare no conflict of interest.

## Author Contributions

N.W. Tam: Designed research, Performed research, Analyzed data, Wrote the paper

R. Dimova: Designed research, Wrote the paper, Other (project supervision)

A. Cipitria: Designed research, Wrote the paper, Other (project supervision)

## Supporting Information

Supplementary figures, including additional vesicle characterization data.

## Notes

https://doi.org/10.17617/3.AMI3GV

### Competing Interest Statement

The authors have declared no competing interest.

### Summary of Updates

Updated text to improve readability, added more references on EVs in tissues

