## Supplemental Information for "Breast cancer cell-derived extracellular vesicles accelerate collagen fibrillogenesis and integrate into the matrix"

### Supplemental Materials

Nicky W. Tam[1]*, Rumiana Dimova[1]*, and Amaia Cipitria [1,2,3]*

[1]Max Planck Institute of Colloids and Interfaces, Science Park Golm, 14476 Potsdam, Germany

[2]Group of Bioengineering in Regeneration and Cancer, Biogipuzkoa Health Research Institute, 20014 San Sebastián, Spain

[3]IKERBASQUE, Basque Foundation for Science, 48009 Bilbao, Spain

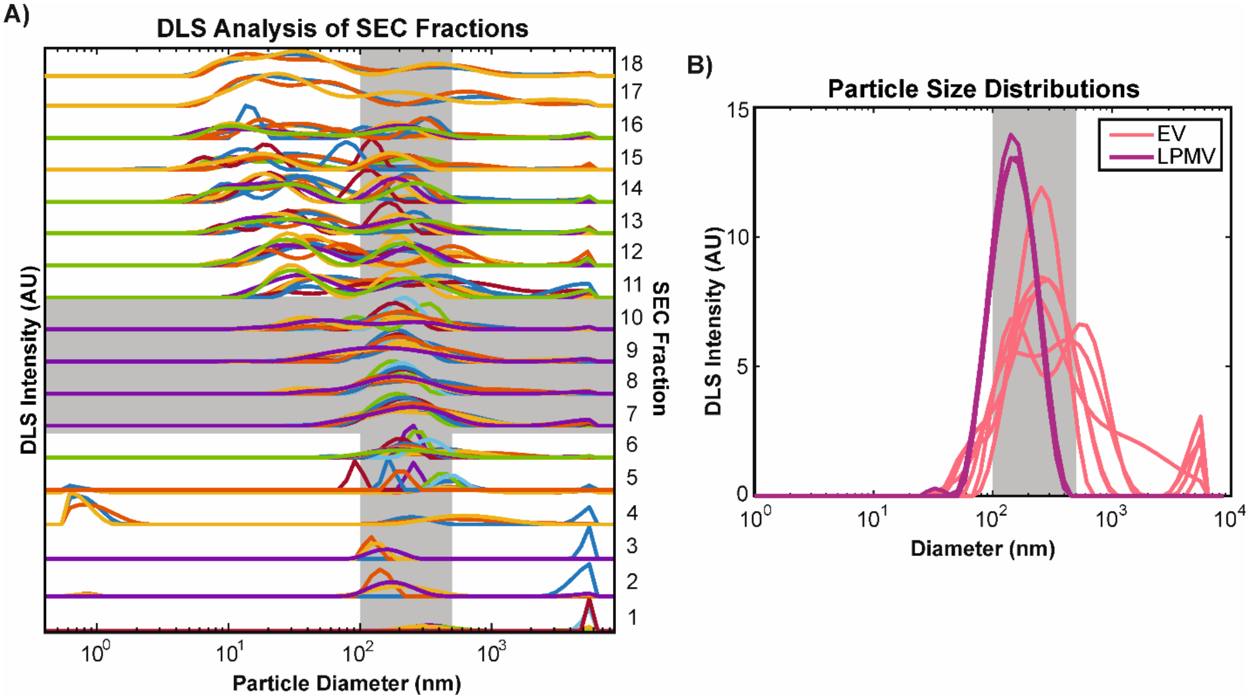


**Supplemental Figure 1**, DLS analysis of EV and LPMV size distribution. Data and figure adapted from the supplemental materials of a previous publication analyzing the same particles.[1] A) Size distributions of particles detected by DLS in the different fractions collected during SEC purification of EVs. Different colours represent different replicates. The target particle size range of 100-400nm and the collected fractions are shaded in grey to show that they intersect. Y-axes of the size distributions represent DLS intensity and have been normalized to show relative enrichment as opposed to absolute abundance. B) Comparison of EV (pink) and LPMV (purple) size distributions. EV traces represent pooled EV fractions. EVs are expected to be more polydisperse because LPMVs are extruded with a defined filter pore size. Different curves represent different replicates.

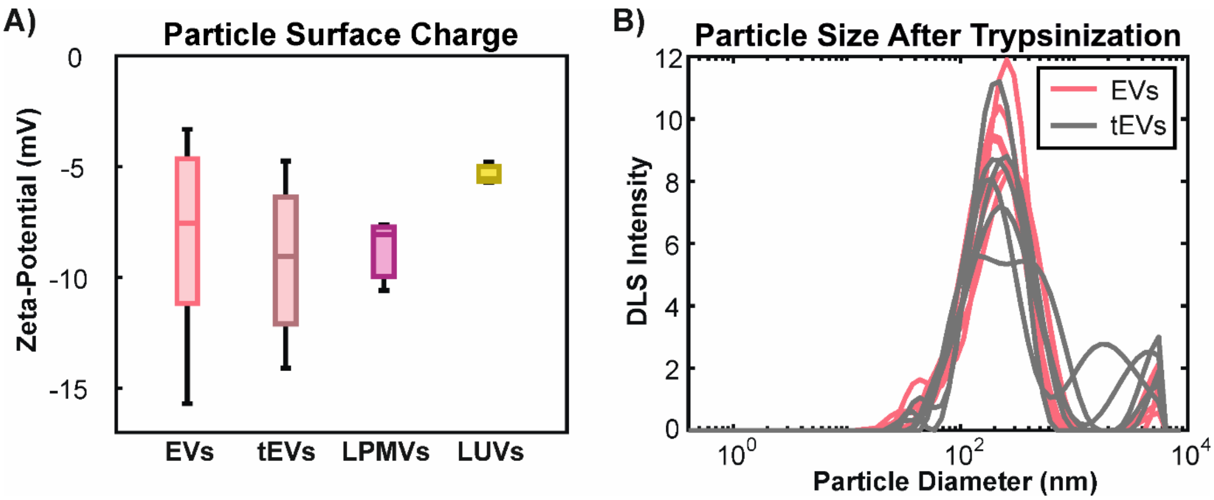


**Supplemental Figure 2**, Effect of trypsinization on EV surface charge and size distribution. Data and figures adapted from the supplemental materials of a previous publication analyzing the same particles.[1] A) Relative surface charges of different particles, as represented by zeta-potential measurements in high ionic strength buffers (HBS). EVs and LPMVs have similar surface charge and trypsinization does not appear to significantly affect EV surface charge. Synthetic DOPC LUVs have a slightly less negative charge. B) Size distributions of EVs before (pink) and after (grey) trypsinization. While overall size does not appear to change much, there appears to be slightly more variability in size and the possible existence of aggregates in tEVs (seen here as extra peaks).

Raw and processed data, as well as MATLAB and Python scripts used for image analysis can be found at the publicly accessible Edmond repository of the Max Planck Society: https://doi.org/10.17617/3.AMI3GV

(1) N.W. Tam, A. Becker, A. Mangiarotti, A. Cipitria, R. Dimova, Extracellular Vesicle Mobility in Collagen I Hydrogels Is Influ-enced by Matrix-Binding Integrins, ACS Nano 18 (2024) 29585–29601. https://doi.org/10.1021/acsnano.4c07186
